# Hypertension is related to a slower radiotracer removal from lateral ventricles

**DOI:** 10.64898/2026.05.07.723657

**Authors:** Katarzyna Olejniczak-Gniadek, Mony J. de Leon, Yi Li, Tracy Butler, Xiuyuan Hugh Wang, Sushruth Manchineella, Chistopher Mardy, Henry Rusinek, Jessica Peña, Yuan Ma, Surendra Maharjan, Liangdong Zhou, Alexus Jones, Emily Tanzi, Silky Pahlajani, Nancy S. Foldi, Thomas Maloney, Carmen Barrios Castellanos, Krista Wartchow, Laura Beth J McIntire, Gloria C Chiang, Lidia Glodzik

**Affiliations:** Brain Health Imaging Institute, Department of Radiology, Weill Cornell Medicine, New York, USA; Neurorehabilitation Clinic, Military Institute of Medicine, Warsaw, Poland; Department of Radiology, NYU Grossman School of Medicine, New York, USA; Dalio Institute of Cardiovascular Imaging, Department of Radiology and Division of Cardiology, Department of Medicine, Weill Cornell Medicine, New York, USA; Department of Epidemiology, Harvard TH Chan School of Public Health, Boston, Massachusetts, USA

**Author notes:** Corresponding author: Lidia Glodzik, Department of Radiology, Weill Cornell Medicine, 407 East 61st Street, New York NY, 10021.

## Abstract

**Background:** Impairment of brain waste removal contributes to Alzheimer’s disease etiology and progression. Although hypertension is a risk factor for dementia, little is known about how it affects measures of clearance in human brain.

**Methods:** Cross-sectional (n=159) and longitudinal (n=94) analysis of the relationship between blood pressure (BP) and brain clearance. The estimate of brain clearance was measured using positron emission tomography (PET) as the rate of radiotracer (MK-6240) efflux from the lateral ventricles in the 10–30-minute window after tracer injection. We also examined cerebral blood flow, PET-derived tau deposition in the medial temporal lobe, cognition and plasma biomarkers of neurodegeneration. At baseline we compared participants with (n=88) and without (n=71) hypertension. For longitudinal analyses we defined two groups based on systolic BP trajectories from baseline to follow-up: as long-term controlled (n=76) or uncontrolled BP (n=18).

**Results:** At baseline, subjects with hypertension had lower ventricular clearance than normotensive controls (Cohen’s d=0.53, p=0.001). Over the course of the observation period (median 1.85 years) subjects in the uncontrolled BP group experienced a steeper reduction in clearance rates (β=-5.88) than subjects in the controlled BP group (β=-0.81, interaction p=0.039).

**Conclusions:** Our study suggests that hypertension impairs brain clearance of fluids.

---

High blood pressure (BP) is a common vascular risk factor in people older than 65 and an established risk factor for dementia including Alzheimer’s disease (AD).^1^ By affecting multiple aspects of the human neurovasculome, hypertension is detrimental to perfusion, cerebral autoregulation, and blood-brain barrier function.^2^ Glymphatic system, where the cerebrospinal fluid (CSF) flows along para-arterial spaces, enters the brain parenchyma, mixes with interstitial fluid and transports waste products away into paravenous spaces,^3^ is another possible target of hypertension. Although there are likely multiple physiological forces driving glymphatic clearance including sleep, circadian rhythms, lifestyle factors^4^ and breathing,^5^ cerebral blood flow (CBF) and arterial pulsations substantially contribute to fluid movement through this system.^6,7^ Consequently, hypertension, leading to changes in brain micro-and macrovasculature,^2^ impairment in CBF and brain autoregulation, has a significant potential to compromise clearance function. Animal studies showed that both acute^7^ and chronic hypertension^8^ impair the flow in paravascular spaces, but little is known whether hypertension affects brain clearance in humans. This gap is partly due to the lack of non-invasive methods to assess CSF circulation in vivo. Human studies using intrathecal contrast injections with subsequent serial MRI scanning recapitulate some aspects of animal glymphatic circulation such as route of tracer entry and effects of sleep deprivation.^9^ However this technique is time-consuming and invasive. A diffusion tensor imaging analysis along perivascular spaces has been proposed as a contrast-free method to assess glymphatic function. Still, the localized nature of measurements and uncertainty about the biological meaning of this marker,^10^ limits its potential to estimate fluid dynamics.

We have previously developed a method for estimating CSF clearance using positron emission tomography (PET) with low molecular weight tracer. CSF clearance is computed directly from the CSF time-activity curves (TAC) in the lateral ventricle (LV).^11^ We validated the method by showing that the rate at which radiotracer is cleared from the ventricle is inversely related to amyloid deposition,^11-13^ AD diagnosis,^11^ cognition,^12^ and age.^13^ Moreover, traumatic brain injury (TBI) uncoupled the correlation between CBF and clearance seen in controls.^14^

In this study we investigated the relationship between ventricular CSF clearance (*kCSF-LV*) and BP. Based on our previous work on hypertension and CBF,^15^ we hypothesized that kCSF*-LV* and CBF will be related and they will be lower in hypertensive participants than in normotensive peers. We examined whether kCSF*-LV* differences persist after accounting for CBF. We tested how BP control over time affects clearance and CBF. Finally, we investigated whether *kCSF-LV* can predict cognitive performance and plasma biomarkers of AD, neurodegeneration and neuroinflammation.

## Methods

### Participants

were recruited at the Brain Health Imaging Institute, Weill Cornell Medicine between 2020 and 2023 for studies investigating effects of hypertension, aging and AD on brain clearance. They were recruited form the community through mailing list and the word of mouth. Exclusion criteria were cortical stroke, brain tumour, neurodegenerative disorders, substance abuse and cognitive impairment. All participants gave written informed consent. All underwent magnetic resonance imaging (MRI) to assess brain CBF, PET using the tau tracer MK-6240 to assess brain clearance, physical and neurological exams, cognitive testing and laboratory assessments. Ninety-four individuals completed follow-up PET examinations and were included in longitudinal analyses. Figure S1 summarizes participants’ flow and reasons for missing follow-up.

### Cognitive status

was determined by a physician’s assessment and the Clinical Dementia Rating (CDR).^16^ Only subjects with CDR=0 were included. Cognitive performance was measured using the Uniform Data Set battery.^17^ The following measures were derived: a) delayed recall: list recall from Rey Auditory verbal learning task (RAVLT) and an average of verbatim and paraphrase Craft story recall, b) working memory (number span backward), c) sematic memory (an average of animal and vegetable fluency), d) learning (total score for RAVLT trials 1-5), e) attention (number span forward), f) executive functions with phonemic fluency (an average of F and L letter fluency). All tests were adjusted for age, sex and education.^17,18^ Z-scores were used for the Craft story, number span and fluency measures and t-scores for the RAVLT.

### Blood pressure

was measured on the left upper arm using a sphygmomanometer with the participant in a sitting position.^19^ Further details are presented in the Supplement. *Hypertension* was defined as current antihypertensive treatment and/or BP≥140/90 mmHg.^1^ The liberal, older threshold of 140/90 was in place when this project was designed. Supplement shows analyses based on current BP guidelines^1^ with a threshold of 130/80.

### Long-term controlled and uncontrolled BP

Participants with baseline and follow-up BP measurements were classified into two groups. The “uncontrolled BP” group consisted of individuals with systolic BP (SBP)≥140 at both visits and those with normal BP at baseline but elevated BP at follow-up. The “controlled BP” group consisted of participants with SBP<140 both at baseline and follow-up and those whose BP was uncontrolled at baseline but controlled at follow-up. The focus on SBP was motivated by greater risk of cardiovascular outcomes associated with high SBP in people older than 50.^20^

#### Medication

We recorded the use of angiotensin receptor blockers, angiotensin converting enzyme inhibitors, beta-blockers, diuretics, calcium channel blockers, and statins.

#### Diabetes mellitus

was defined as fasting glucose ≥ of 126 mg/dL, or hemoglobin A1c ≥ 6.5%,^21^ or current use of glucose lowering medication.

Blood samples were tested for complete blood count, liver function, metabolic, and *lipid profile. Body mass index (BMI)* was calculated as weight/height^2^ [kg/m^-2^].

#### Smoking

status was defined as positive if the participant was a current smoker or smoked within last 10 years.

### MR imaging

was performed on research dedicated 3T Prisma system (Siemens, Erlangen, Germany) and consisted of T1- and T2-weighted imaging, Fluid-Attenuated Inversion Recovery (FLAIR), multi-echo gradient echo T2 and pseudo-continuous arterial spin labeling (pCASL) sequences (parameters given in the Supplement).

#### Regions of interest

for PET and CBF analyses were segmented using Freesurfer (FS) version 7.1.22 T1 and T2 images were used to increase quality of FS parcellation.

#### CBF mapping

was performed using ASLtbx.^23^ First, pCASL images were motion corrected: volume frames were realigned to the mean of all volumes using MCFLIRT. Second, we performed a voxel-wise subtraction of label and control images, followed by averaging of all the subtracted perfusion weighted images. Perfusion weighted and M0 images were then fed into the ASLtbx for CBF calculation. Calculated CBF was coregistered to subjects’ T1 images in FS atlas space. Subsequently, CBF values were extracted from the entire neocortex obtained through FS segmentation.

Periventricular and deep *white matter hyperintensities* (WMH) were graded separately on FLAIR images using 0-3 Fazekas scale.^24^

#### Microbleeds

defined on susceptibility weighted images as hypointense lesions 2-10 mm in diameter,^25^ were coded as ‘present’ or ‘absent’.

### PET imaging

was performed on a Siemens Biograph mCT−S (64) slice PET/CT. 18F-MK240 was provided by the manufacturer (Cerveau). After intravenous bolus injection of 185 MBq of ^18^F-MK6240, dynamic brain PET data were acquired in 3-dimensional list mode from 0 to 60 min post-injection. The list-mode data were rebinned and reconstructed into a 400×400×109 matrix with 1×1×2 mm voxels and 26 dynamic frames (12×10 s, 3×1 min, 11×5 min), providing high temporal resolution necessary to capture the early dynamics of tracer delivery and outflow from the lateral ventricles. Image reconstruction was performed with standard Siemens neuro-PET protocols using an iterative TOF-OSEM algorithm.

#### Assessment of CSF ventricular clearance

was carried similarly to previously described.^11-13^ The dynamic frames were re-aligned to the image formed by the summation of all frames acquired between 6-30 minutes. The summation image was coregistered to the T1 in FS space using normalized mutual information cost function and the coregistration transformation matrix was saved. The transformation matrix was then applied to all the dynamic frames to align PET data to FS space. To reduce partial volume contamination by brain tissue, LV region was eroded by 3 voxels. Choroid plexus was excluded. For consistency, the transformation matrix was calculated both for baseline and follow-up PET to the baseline MRI, then the same LV region was applied. This ventricular region is presented as ventricular volume in Tables. The ventricular CSF clearance rate (*kCSF-LV*) was calculated by estimating the TAC slope in the LV during 10-30 min period. This time frame reduces the potential contribution of blood based counts in the choroid plexus.^12^ *kCSF-LV* was inversely related to ventricular volume, thus the volume was used as a covariate in the analyses.

#### Assessment of tau deposition

was based on 18F-MK240 standardized uptake value ratio (SUVR) calculated using a summed image during 40 to 60 min. Cerebellar cortex served as a reference region. This timeframe was used due to the requirement of scanning dynamically for 60 minutes immediately after injection to achieve the main study goal and limited feasibility of extending scan beyond this period or rescanning in the recommended later time window.^26^ We focused on medial temporal lobe (MTL), a site of early tau accumulation because our participants had normal cognition and cortical deposition was not expected. Tau status visual reading (tau+/-) was performed using previously defined criteria.^27^ Importantly, they were established on a later (60-90 min) acquisition window.

### Plasma biomarkers

Plasma biomarkers were examined for 72 participants at baseline and 56 at follow-up. Blood was drawn at the time of clinical labs, spun, aliquoted and frozen at -80C until further analyses. Amyloid β42 (Aβ42), amyloid β40 (Aβ40), p-tau217, neurofilament light (NFL), and glial fibrillary acid protein (GFAP) were assessed using Quanterix Simoa HDX technology following the manufacturer’s protocols. Analyses were performed with ALZpath p-Tau217 Advantage Plus and Simoa N4PE Advantage Plus Reagent Kits, for p-tau217 and Aβ42, Aβ40, NFL, GFAP, respectively. In all analyses a Aβ42/40 ratio was used.

### Statistical Analyses

Variables were compared using Chi-squared, Fisher’s exact and Wilcoxon rank sum tests.

Bivariate correlations between *k_CSF-LV_*, CBF, cognitive tests and biomarkers were assessed using Spearman’s rank correlation coefficient. Significant correlations were further tested with generalized linear models (GLM), adjusting for appropriate covariates. Unstandardized β values and 95% confidence intervals (CI) are reported. To test whether between normo- and hypertension,associations between variables differ by groups, interaction terms were added. Group differences were tested with non-parametric Quade ANCOVA, since linear model assumptions were not met. Cohen’s d was calculated using partial eta squared (η2*p*). First, η^2^*p* was converted to Cohen’s *f*:

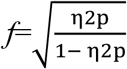

and d was calculated as d=2*f*.

While testing differences in *k_CSF-LV_* between normo- and hypertension, we adjusted for ventricular volume, CBF, age, sex, diabetes and statin use. When testing plasma biomarkers, age and glomerular filtration rate (GFR) were covariates.

CBF values were missing at baseline for 2 out of 159 subjects, and while comparing hyper-and normotensive subjects, these 2 values were imputed based on equation derived from our sample:

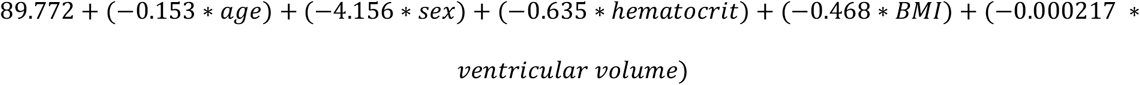

To examine whether variables changed between visit 1 and 2, we used generalized estimating equations (GEE). The variables of interest were dependent, and age, sex and visit were covariates. To test the influence of BP control on variables changes, we added group (i.e. BP uncontrolled vs. BP controlled), and *group*visit* interaction.

Then, with GEE, we tested how SBP changes (independent variable) were related to changes in other variables. Next, we examined whether the relationships between SBP changes and changes in other variables were different based on longitudinal BP control status. For this purpose, in GEE models SBP, group based on BP control, and *group*SBP* interaction were predictors.

Finally, we examined whether changes in *k_CSF-LV_* or CBF predicted changes in MTL tau, plasma biomarkers or cognition. Separate GEE models were employed for each pair of variables with clearance or CBF as predictors. To test whether associations between variables differ by groups interaction terms were added to GEE models.

A p-value<0.05 was deemed significant. For analysis of cognitive tests p-value was set at 0.05/7=0.007. For the analysis of plasma biomarkers, it was set at 0.05/4=0.0125. Tables were created using *gtsummary* package in R. SPSS (version 28, SPSS, Inc., Chicago, IL) software was used for all other analyses.

## Results

### Baseline analyses

The study included 159 participants with median age of 70.1 (interquartile range IQR 11.8), education 17 years (IQR 2), 83 (52%) were women. Approximately 78 % of participants were White, 17% Black, 3% Asians and 2% of other races. Hypertension at baseline was present in 88 participants.

### Entire group

#### Relationships between k_CSF-LV_ and CBF

In the entire group *k_CSF-LV_* and CBF were positively related (GLM CBF β=0.486, 95% CI 0.272-0.699, p<0.001, adjusted for ventricular volume).

#### Relationships between k_CSF-LV_ or CBF and cognitive performance, tau deposition and plasma biomarkers

No associations were found.

### Hypertension vs. Normotension

The two groups differed in age, sex distribution, hemoglobin A1c, statin use, lipid profile, BMI, C-reactive protein, GFR, ventricular volume, cortical CBF, WMH burden (Tab. 1 and Tab.S1).

**Table 1.**
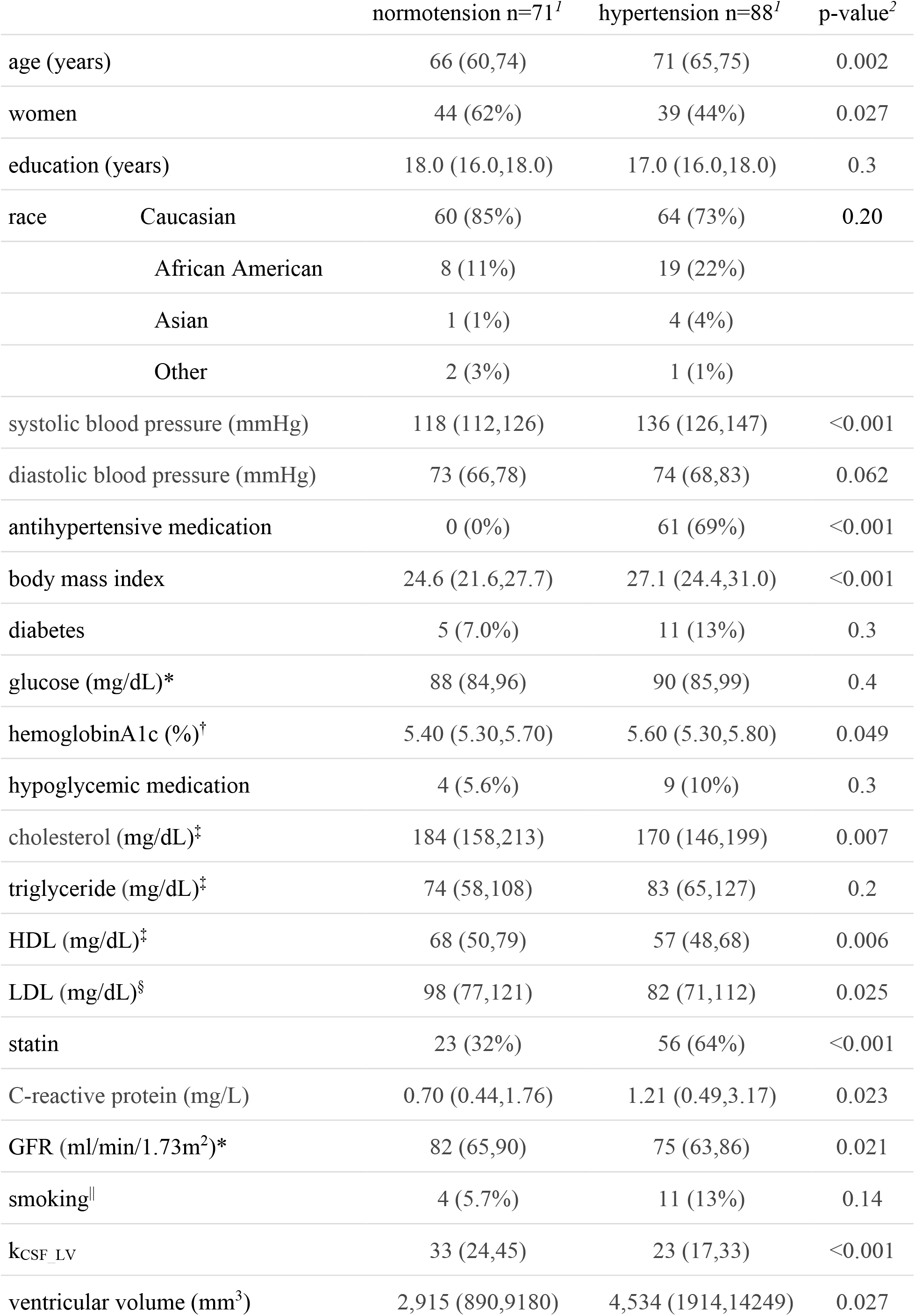
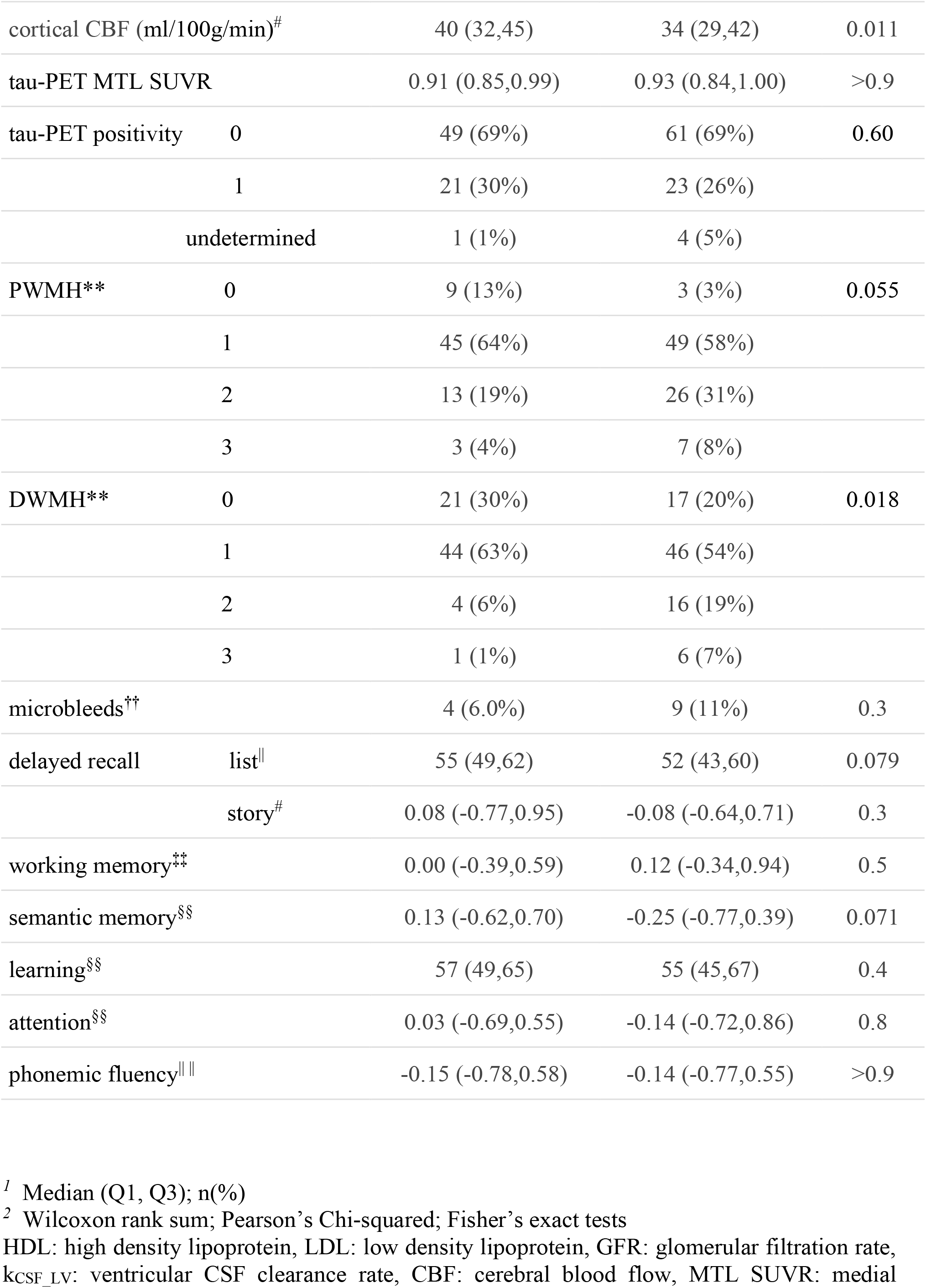
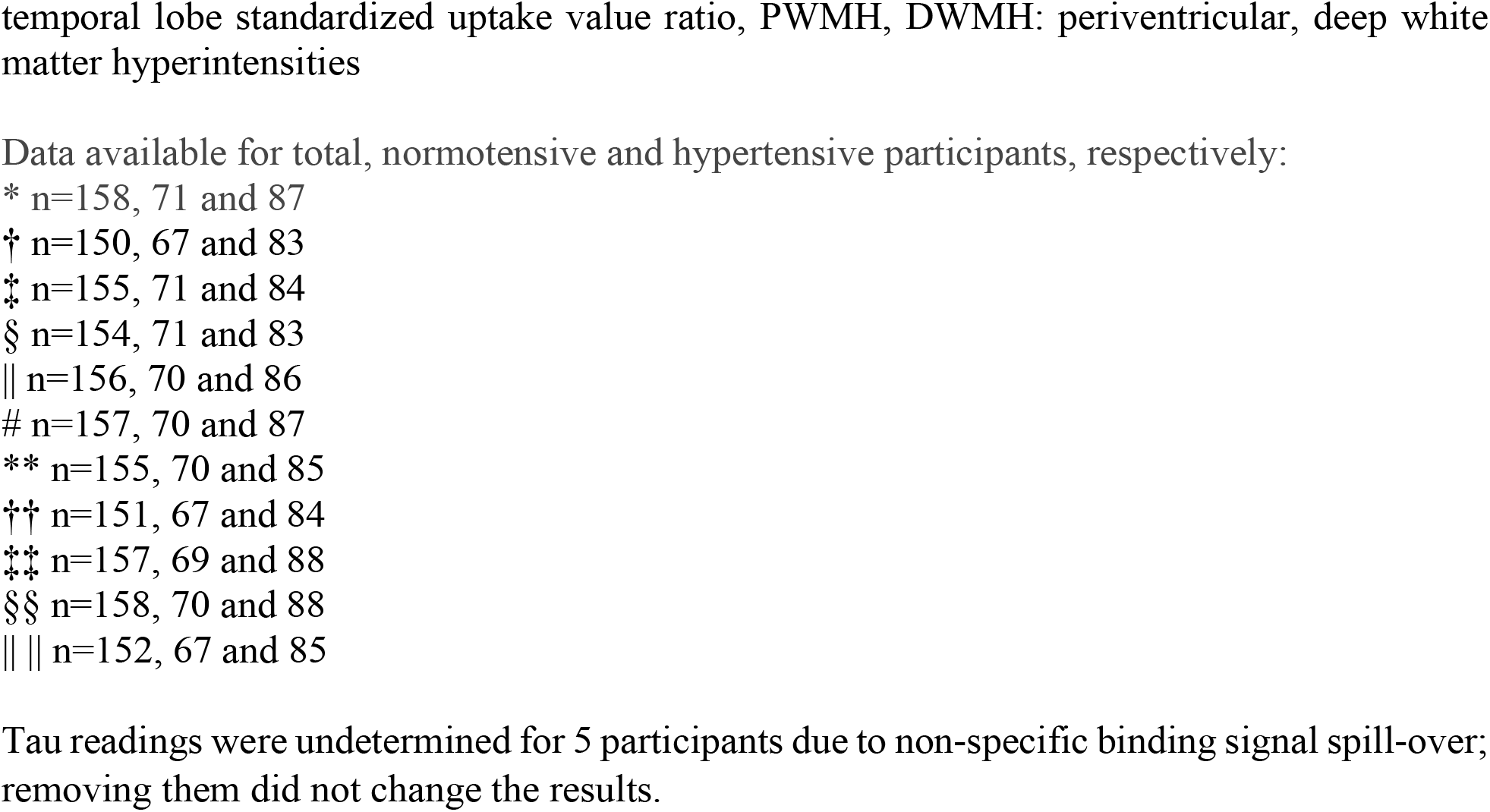
Baseline characteristics of the study group.

Hypertensive participants had lower *k_CSF-LV_* than normotensive controls: Quade ANCOVA adjusted for ventricular volume, CBF, age, sex, diabetes, and statin use F1,158=11.06, p=0.001, Cohen’s d=0.53.

#### Effects of hypertension on relationships between k_CSF-LV_ and CBF

The associations were similar in both groups.

#### Effects of hypertension on relationships between k_CSF-LV_ or CBF and cognitive performance, tau deposition and plasma biomarkers

No associations were found in either group.

### Longitudinal analyses

Follow-up (median 1.85 years, IQR 0.62) was available for 94 subjects. Participants studied only once (n=65) were more likely to be White (89% vs 70%, exact test=10.14, p=0.009) and performed better on RAVLT learning task (p=0.045). No other differences were found when comparing variables listed in Tab. 1. Tab.S2-S3 describe longitudinally studied participants, stratified by baseline hypertension status.

### Entire group

#### Changes in variables over time

GEE models with age, sex and visit as predictors showed that neither changes in *k_CSF-LV_* (β=-1.79, p=0.12), nor in CBF (β=-1.37, p=0.07) reached significance. The results for *k_CSF-LV_* remained similar after adding CBF changes to the model (p=0.06).

MTL tau increased (β=0.015, p=0.04). After excluding subjects with undetermined tau visual tau assessment, the results were no longer significant, p=0.12.

Delayed list recall (β=3.7, p=0.001), learning (β=6.2, p<0.001), semantic memory (β=0.29, p<0.001), and phonemic fluency (β=0.24, p=0.006) improved. GFAP (β=10.3, p=0.003) and NFL (β=2.78, p=0.01) increased, while Aβ42/40 ratio (β=-0.005, p<0.001) decreased.

#### Relationship between k_CSF-LV_ and CBF

There was no association between baseline CBF and the rate of change in *k_CSF-LV_*. GEE analysis with CBF changes as a predictor of *k_CSF-LV_* changes showed that CBF increases were related to *k_CSF-LV_* increase (β= 0.32, p=0.007).

#### Relationships between changes in k_CSF-LV_ or CBF and changes in cognitive performance, tau deposition and plasma biomarkers

GEE analyses with *k_CSF-LV_* or CBF changes as a predictor did not reveal any associations with changes in cognition, tau or fluid biomarkers.

#### Relationships between changes in SBP and changes in k_CSF-LV_, CBF, cognitive performance, tau deposition and plasma biomarkers

The SBP increase was related to reduction in *k_CSF-LV_* (β=-0.16, p=0.004). The results remained the same after adding CBF to the model (SBP p=0.003). Changes in SBP were not related to changes in CBF, MTL tau, cognitive tests or fluid biomarkers.

### Long -term controlled vs. uncontrolled BP

Participants with long-term controlled (n=76) and uncontrolled (n=18) BP are presented in Tab.S4-S5 (baseline) and Tab.S6-S7 (follow-up). Participants with uncontrolled BP had lower baseline GFR, higher p-tau217 and higher NFL. Adjustments for age and GFR, did not affect plasma results.

#### Changes in variables over time

GEE analyses with age, sex, visit, BP control status, and *BP control*visit* interaction as predictors showed that in the group with uncontrolled BP *k_CSF-LV_* decreased at a greater rate than in the controlled BP group (interaction p=0.039, uncontrolled group β=-5.88, controlled group β=-0.81, Fig. 1a). Trajectories of CBF did not differ between the two groups (Fig. 1b). The results for *k_CSF-LV_* remained similar after adding longitudinal CBF to the model (interaction p=0.07). CBF itself did not contribute to the model, and missing data reduced statistical power.

**Figure 1.**
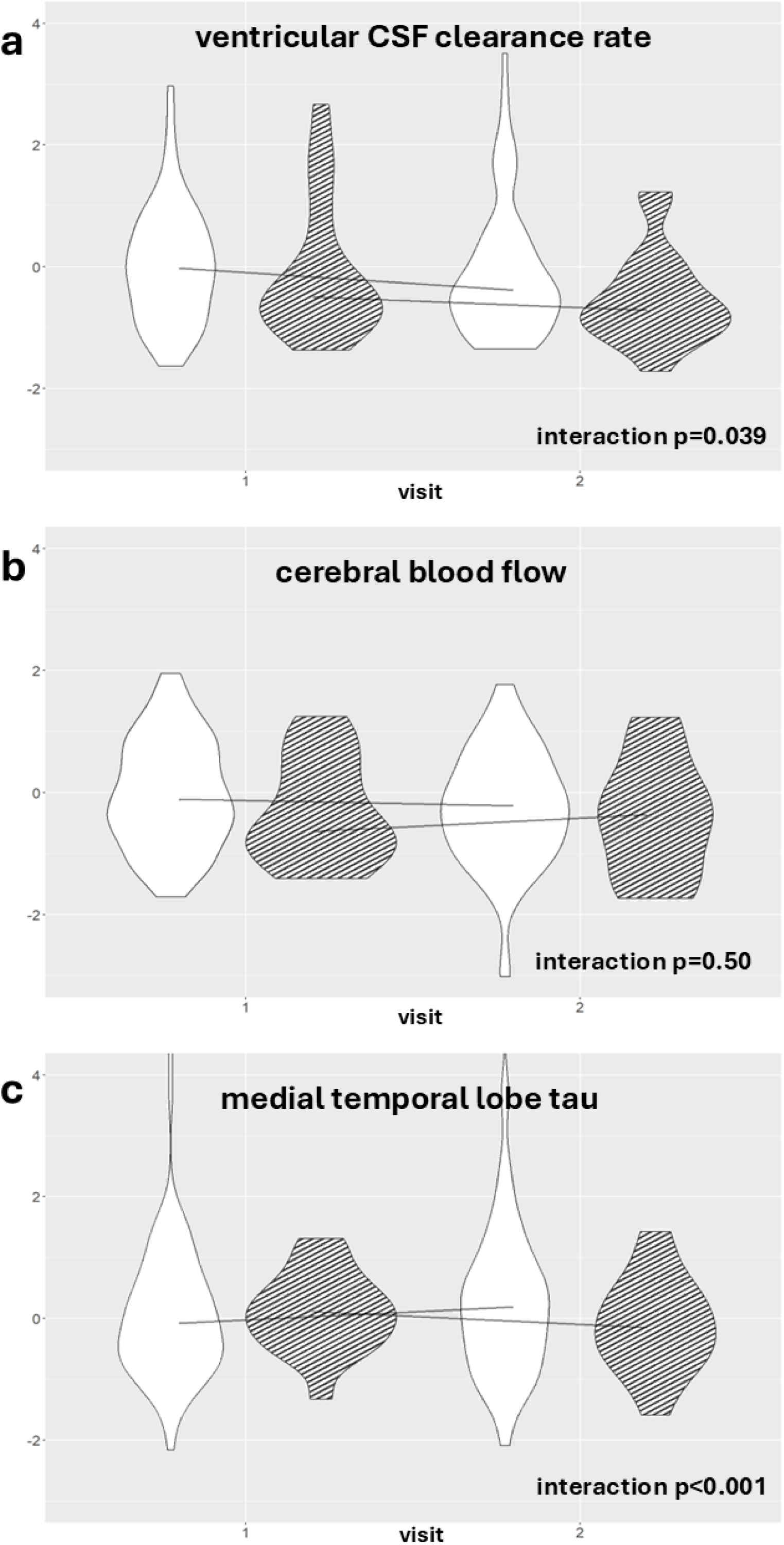
Trajectories of main variables in longitudinally controlled (n=76, white) and uncontrolled BP (n=18, stripes). P-values come from GEE models with age, sex, visit, BP control group and *group*visit* interaction. There were no between-group differences at any given time-point. All values were z-scored, lines connect the medians.

There was a significant *BP control*visit* interaction (p<0.001) for MTL tau, such that the greater MTL tau accumulation was observed in subjects with controlled BP (β=0.024, uncontrolled group β=-0.024, Fig. 1c). Results remained the same when subjects with undetermined visual tau assessment were excluded.

Trajectories of plasma biomarkers (Fig. 2a-d) and cognitive performance or associations between *k_CSF-LV_* and CBF did not differ between participants with controlled and uncontrolled BP.

**Figure 2.**
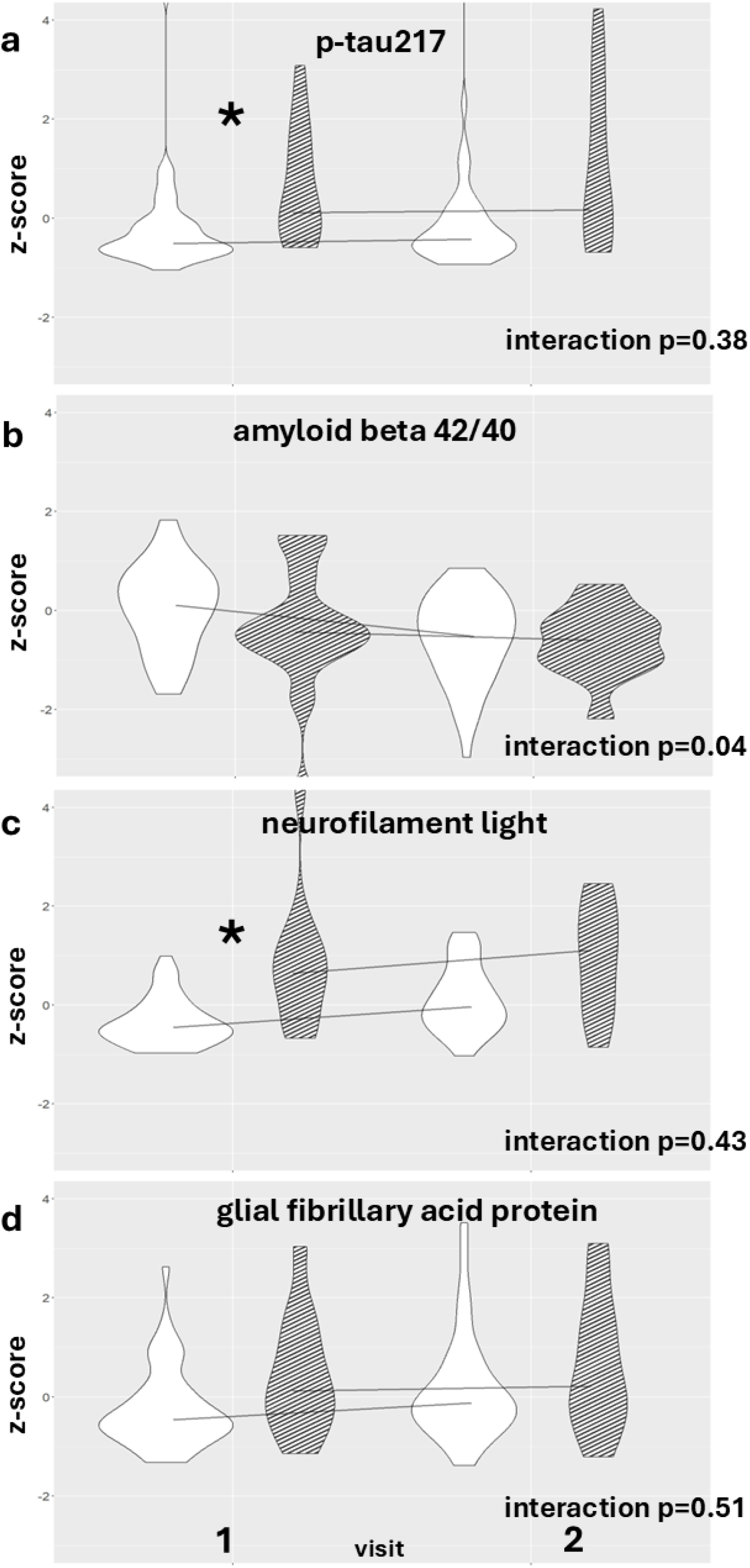
Trajectories of plasma biomarkers in longitudinally controlled (n=76, white) and uncontrolled BP (n=18, stripes). P-values come from GEE models with age, sex, visit, BP control group and *group*visit* interaction. None of the *group*visit* interactions were significant at p<0.0125. Asterix denotes between-group differences at a given time-point at p<0.0125 (age and GFR adjusted). All values were z-scored, lines connect the medians.

#### Relationships between changes in SBP and changes in other variables

GEE analyses were used with SBP, BP control status and their interaction as predictors. Subjects with persistently uncontrolled BP had a more negative SBP–k_CSF-LV_ slope than subjects with controlled BP (interaction p=0.04, uncontrolled group β=-0.37, controlled group β=-0.10). The results remained similar after adding CBF to the models (interaction p=0.095).

Changes in SBP were not related to changes in CBF, MTL tau, cognitive tests or fluid biomarkers in either group except for learning. There was an inverse relationship between SBP and learning in the controlled group but not in the uncontrolled one (interaction p=0.008, controlled group β=-0.13, uncontrolled group β=0.22).

#### Relationships between changes in k_CSF-LV_ or CBF and cognitive performance, tau deposition and plasma biomarkers

Changes in *k_CSF-LV_* or CBF were not associated with changes in cognition, tau or fluid biomarkers in either group.

### Hypertension based on 130/80 threshold

When 130/80 threshold was used, hypertensive patients still showed lower clearance than normotensive peers (Supplemental material). Longitudinally, the uncontrolled group (SBP>130mmHg) showed steeper reduction in *k_CSF-LV_* than subjects with controlled BP and stronger relationship between changes in SBP and clearance changes. However, these differences were attenuated after adding CBF to the models.

## Discussion

We show that the rate at which intravenously injected tracer was cleared from lateral ventricles was lower in participants with hypertension. The effect size was moderate. Moreover, the longitudinal dynamic of this metric differed based on long-term BP control. Specifically, participants with uncontrolled hypertension showed not only lower clearance at baseline but also greater reduction in clearance over time. Lastly, clearance reduction was associated with SBP increase and this effect was driven by the group with uncontrolled BP. These results indicate that elevated BP impairs brain fluid clearance and suggests another pathway linking hypertension to neurodegeneration.

Pertinent to our findings, in a previous animal study, a rapid reduction of BP resulted in a faster removal of dye injected into the cisterna magna.^28^ The efflux route lead through lymphatic vessels. The outflow path from the ventricle in our study is unclear, but suggested pathways include, among others, passage from the ventricles to subarachnoid and para-arterial spaces, subsequent mixing with interstitial fluid, and exit through the veins or through meningeal lymphatic and arachnoid granulations.^9^ In the above study by Jukkola^28^ the driving force pushing the CSF out of the brain was increase in the intracranial pressure caused by vessels dilation and blood inflow. Although, this explanation is more likely in an acute experiment, persistent vasoconstriction from to high BP could have been a cause of reduced tracer removal also in our study. Stiffening and remodelling of vessels from long-term high BP is another explanation in clinical settings.

In accordance with our initial hypothesis both at baseline and longitudinally *k_CSF-LV_* was associated with CBF. This may suggest the importance of CBF in the hypothetical outflow route through paravascular spaces or indicate more efficient initial tracer delivery to the ventricle. Irrespective of the nature of the relationship between k_CSF-LV_ and CBF, the association between k_CSF-LV_ and BP control held both cross-sectionally and longitudinally even after accounting for CBF. Moreover, longitudinal CBF changes were not different in groups with controlled and uncontrolled BP. In all, this suggests that the effect of BP on clearance is not exerted merely through CBF. Opposite to our previous findings in TBI patients,^14^ hypertension did not result in decoupling of the k_CSF-LV_-CBF association. Possibly, this phenomenon occurs with more acute or severe injury.

Longitudinal changes in k_CSF-LV_, CBF and MTL tau in the entire group showed expected trends. Plasma biomarkers became more abnormal with time. These results are congruent with previous observations showing longitudinal increases in NFL and GFAP, and aβ42/40 ratio reduction in a community based cohort.^29^ Likely, GFAP and NFL tracked senescence and neuronal damage.^30^ Cognitive performance of our participants improved. This finding can be explained by expected learning effects^31^ in cognitively healthy subjects. Apparently higher levels of pathology are necessary to abrogate learning effects, and a longer time is needed for cognitive decline to manifest.

In the group with persistently uncontrolled BP p-tau217 and NFL were higher already at baseline. It suggests that even in cognitively normal individuals, uncontrolled hypertension is associated with plasma biomarkers of neurodegeneration. Similarly, earlier studies showed associations between higher p-tau217, GFAP and NFL and vascular stiffness^32^ and others found that PET-amyloid negative subjects with high plasm a p-tau217 were more likely to have hypertension than subjects with low p-tau.^33^ Although these differences can result from renal impairment, in our case, the uncontrolled group had higher values even after adjusting for age and GFR. Notably, despite baseline differences, trajectories of plasma biomarkers did not differ while clearance worsened at a higher pace in the uncontrolled group. Arguably, p-tau217 and NFL reached severity thresholds faster, k_CSF-LV_ is a better dynamic indicator of hypertensive pathology or smaller sample size for plasma analytes prevented us from seeing significant differences.

Surprisingly MTL tau increased only in the controlled BP group. In an earlier study, a sharp mid-life decline in diastolic BP was associated with more entorhinal tau deposition approximately 16 years later.^34^ Despite much longer follow-up time that is congruent with our findings. Since our subjects were on average a few decades older, it could put them at higher risk for hypoperfusion and tau accumulation in a shorter time. These observations also emphasize that BP control in older individuals is a balancing act, with uncontrolled and too-tightly controlled BP both potentially detrimental to brain health. Our prior study similarly showed that in older individuals with hypertension, aggressive BP control may not mitigate its negative effects on the brain.^15^

Neither at baseline nor longitudinally did we find evidence that clearance or CBF were related to plasma biomarkers or cognition. Possibly, some other mechanisms play more important role. Alternatively, CBF and clearance affect them through intermediary mechanisms which were not captured in this study.

### Limitations

The main limitation of study is the relatively unconventional method of looking at brain clearance. Although our initial observation of clearance reduction in AD using PET derived method^11^ was confirmed in an independent group,^35^ it remains unclear what aspects of CSF efflux are captured by our measure. It is rather general estimate of outflow through multiple possible paths: subarachnoid spaces to paravascular channels, lymphatic vessels, arachnoid granulations, and nasal turbinates.^9,11^ Contribution of each route remains unknown.

We have used an outdated threshold of 140 mmHg. Analyses based on the threshold of 130 mmHg showed that groups defined by higher threshold had more pronounced difference in clearance and stronger SBP-clearance association than groups established using 130 mmHg.

This suggests more abnormalities at higher BP. Still, we do not know whether this relationship is truly causal or only correlative.

Due to the requirement of scanning immediately after injection to measure CSF clearance, we derived tau deposition from the 40–60-minute frame. This is earlier than the 90–120-minute window recommended for tau assessment. Mitigating against this limitation, equilibrium is generally reached within 60 minutes except in individuals with later stage AD and high tau burden.^26^ Since our subjects were cognitively healthy, high level of tau accumulation were not expected. In addition, we have recently shown that SUVR derived from 40-60 and 90-120 windows are highly concordant (Hu&Li, in review).

Plasma biomarkers were not assayed in all subjects, and this might have influenced our assessment of relationships between biomarkers and clearance.

### Perspective

We show that the rate of low-molecular weight radiotracer removal from the lateral ventricle in the first 30 minutes after injection is significantly lower in hypertensive participants. Moreover, this process declines at a steeper pace in subjects with persistently high BP, as compared to those in with SBP<140mmHg. Although more human studies are necessary to confirm relationships between measures of brain fluid circulation and hypertension, our study indicates that, similarly as in animal experiments, high BP impairs brain clearance of fluids. Even though exact mechanisms remain unknown, this finding further suggests the benefits of treatment.

## Novelty and Relevance What Is New

This is a study using PET-based method to assess brain clearance in participants with and without hypertension.

### What Is Relevant

Hypertension was associated with lower rate of tracer efflux from lateral ventricles. Participants with poor long-term BP control showed steeper decline in clearance rates than the group with controlled hypertension.

### Clinical/Pathophysiological Implications

Although more human studies are necessary to confirm relationships between cerebral fluid circulation and hypertension, our study suggests that high BP impairs brain clearance and underscores the importance of BP control.

